# Atomistic ensemble of active SHP2 phosphatase

**DOI:** 10.1101/2023.05.04.539460

**Authors:** Massimiliano Anselmi, Jochen S. Hub

**Affiliations:** Theoretical Physics and Center for Biophysics, Saarland University, 66123 Saarbrücken, Germany

## Abstract

SHP2 phosphatase plays an important role in regulating several intracellular signaling pathways. Pathogenic mutations of SHP2 cause developmental disorders and are linked to hematological malignancies and cancer. SHP2 comprises two tandemly-arranged SH2 domains, a catalytic PTP domain, and a disordered C-terminal tail. Under physiological, non-stimulating conditions, the catalytic site of PTP is occluded by the N-SH2 domain, so that the basal activity of SHP2 is low. Whereas the autoinhibited structure of SHP2 has been known for two decades, its active, open structure still represents a conundrum. Since the oncogenic mutant SHP2^E76K^ almost completely populates the active, open state, this mutant has been extensively studied as a model for activated SHP2. By molecular dynamics simulations and accurate explicit-solvent SAXS curve predictions, we present the heterogeneous atomistic ensemble of constitutively active SHP2^E76K^ in solution, encompassing a set of conformational arrangements and radii of gyration in agreement with experimental SAXS data.

## INTRODUCTION

Src-homology-2–containing protein tyrosine phosphatase 2 (SHP2), encoded by the *PTPN11* gene, is a classical non-receptor protein tyrosine phosphatase (PTP).^1^ SHP2 has emerged as a key downstream regulator of several receptor tyrosine kinases and cytokine receptors, functioning as a positive or negative modulator in multiple signaling pathways,^2, 3^ such as the Ras/mitogen-activated protein kinase (MAPK) pathway, in the context of which, SHP2 acts as a positive transducer of proliferative and antiapoptotic signals.^4^ Germline mutations in the human *PTPN11* gene have been associated with Noonan syndrome and with Noonan syndrome with multiple lentigines (formerly known as LEOPARD syndrome), two multisystem developmental disorders.^5–16^ Somatic *PTPN11* mutations were linked to several types of human malignancies,^17–20^ such as myeloid leukemia.^6, 14, 21–24^ SHP2 is nowadays a key target for anti-cancer treatment against drug-resistant metastatic tumors.^25–32^

The structure of SHP2 includes two tandemly-arranged Src-homology-2 domains (SH2), called N-SH2 and C-SH2, followed by the catalytic PTP domain, and a C-terminal tail (Figure 1a-b).^33^ The SH2 domains are structurally conserved recognition elements that allow SHP2 to bind signaling peptides containing a phosphorylated tyrosine (pY).^34^ As the catalytic core of SHP2 shares high homology to those of SHP1 and other PTPs, which play negative roles in cell signaling,^35^ drug design needs to exploit structural features unique to SHP2. It turns out that structural and mechanistic understanding of SHP2 activation is required for designing a new generation of selective inhibitors.

**Figure 1.**
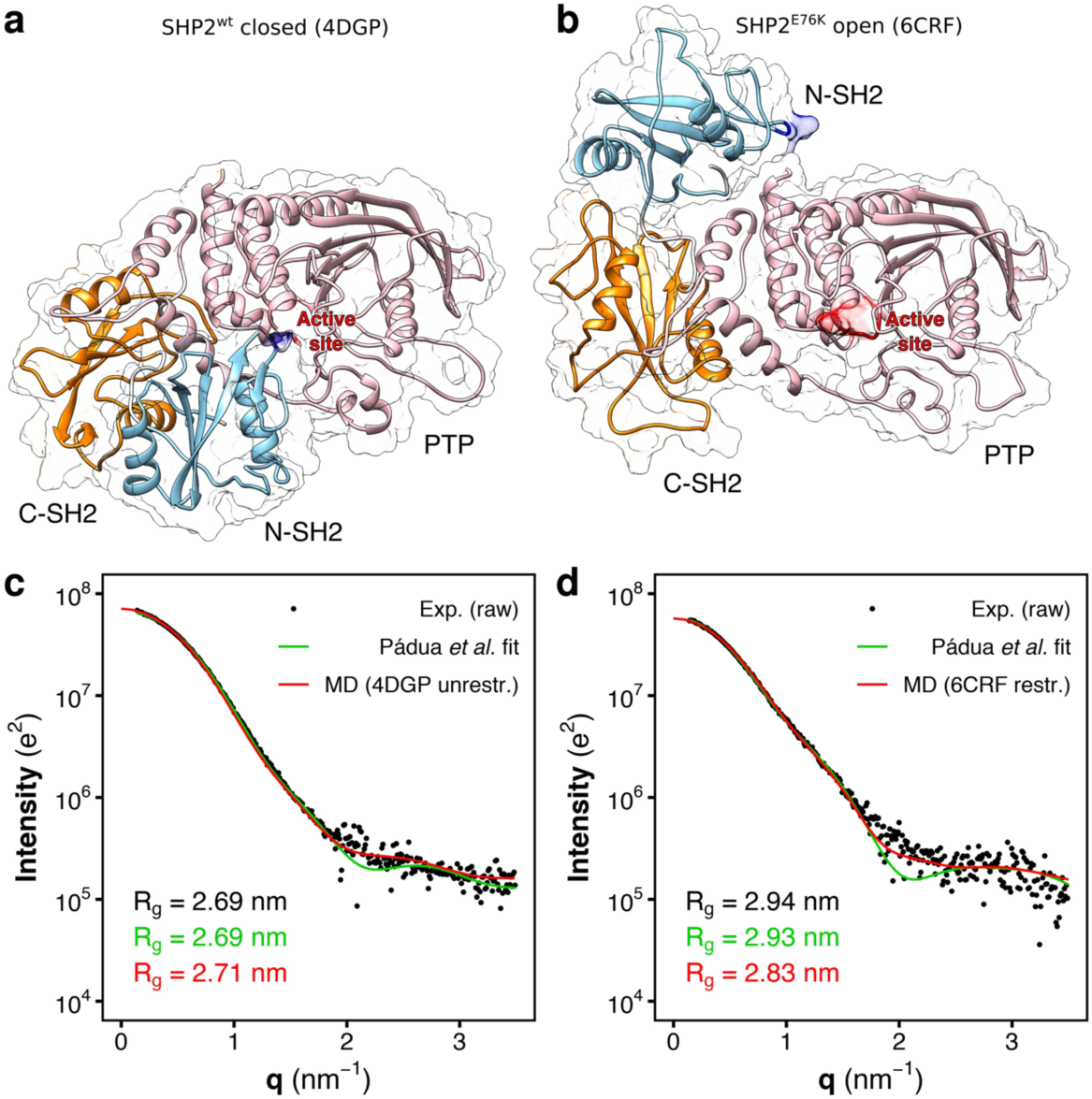
Crystal structures and SAXS curves from closed SHP2 and open SHP2^E76K^. a) Crystal structure of autoinhibited wild-type SHP2 (SHP2^wt^). In the autoinhibited state (PDB ID 4DGP^15^), SHP2^wt^ adopts a closed conformation; the N-SH2 domain (cyan cartoon) blocks the catalytic site (red) of the PTP domain (pink) with the blocking loop (blue). The N-SH2 domain is connected to PTP in tandem with the homologous C-SH2 domain (orange). b) Crystal structure of the constitutively active SHP2^E76K^ mutant. In the active state, SHP2^E76K^ adopts an open conformation with the catalytic pocket exposed to the solvent. The crystal structure of SHP2^E76K^ (PDB ID 6CRF^44^) reveals a 120° rotation of the C-SH2 domain and the relocation of the N-SH2 domain to a PTP surface opposite the catalytic site. c) Small-angle X-ray scattering (SAXS) curve calculated from the solution ensemble of the autoinhibited wild-type SHP2 (unrestrained MD simulation from PDB ID 4DGP,^15^ red line) is compared with the experimental curve (black dots),^46^ reported as raw data, and with the single-structure model fitting by Pádua *et al*. (green line).^46^ d) The SAXS curve calculated from the crystal structure of the constitutively active SHP2^E76K^ mutant (restrained MD simulation from PDB ID 6CRF,^44^ red line) is compared with the experimental curve (black dots),^46^ reported as raw data, and with the single-structure model fitting by Pádua *et al*. (green line),^46^ Experimental, model-estimated, and calculated radii of gyration (*R*_g_) are reported in each panel in black, green, and red font, respectively.

Crystal structures have revealed an allosteric regulation of SHP2 activity.^36^ Under basal conditions, SHP2 is in an autoinhibited state where the N-SH2 domain occludes the catalytic site of the PTP domain (Figure 1a). Association of SHP2 to its binding partners via the SH2 domains favors the release of the autoinhibitory N-SH2–PTP interactions, rendering the catalytic site accessible to substrates.^36–40^ The basal activity of wild-type SHP2 is low,^8^ suggesting that under non-stimulating conditions the protein is almost completely in the autoinhibited, closed state and the fraction of open structure is virtually negligible.

Critically, in presence of a saturating concentration of monophosphorylated activating peptides the activity of wild-type SHP2 increases only four- to fivefold;^8^ hence, the binding of a single phosphopeptide to the N-SH2 domain triggers a mild activation, suggesting that the N-SH2 domain remains in the neighborhood of the catalytic site.^8, 41, 42^ Bidentate phosphopeptides, such as BTAM (bisphosphoryl tyrosine-based activation motif), contain two pY residues and can potentially bind simultaneously to the N-SH2 and C-SH2 domains. BTAMs are stronger activators than monophosphopeptides;^8^ in fact only the binding of another phosphopeptide to C-SH2, connected to the first phosphopeptide by a linker of at least 40 Å length,^41^ drives to the complete dissociation of the N-SH2 domain from the catalytic site and, thereby, to further activation of SHP2 by 10 to 20 folds relative to basal activity.^36, 42^ However, the activity of wild-type SHP2 does by far not reach the maximum activity of the truncated construct of SHP2 without N-SH2 (ΔN-SH2),^43^ indicating that, even under stimulating conditions, a significant fraction of wild-type SHP2 retains the autoinhibited, closed conformation.

The factors influencing the equilibrium between the autoinhibited and the active state of SHP2 have been extensively studied by mutagenesis experiments.^6, 8^ Many pathogenic mutations of SHP2 cluster at the N-SH2/PTP interface, where they destabilize the N-SH2/PTP interactions, causing constitutive activation of SHP2.^6^ Amongst these mutations, the basally active, leukemia-associated, E76K mutant is particularly relevant and extensively studied as a reference system. Since the Glu^76^ side chain forms a salt-bridge with Arg^265^ of the PTP domain,^36^ the E76K substitution replaces the salt-bridge with an unfavorable electrostatic and steric interaction between Lys^76^ and Arg^265^. As a consequence, among the pathogenic mutants, E76K displays the highest levels of basal activity, indicating that SHP2^E76K^ almost completely populates the active, open state, even under non-stimulating conditions.^6, 8, 43^

For that reason, SHP2^E76K^ has been often taken as a model system for the open state of full-length SHP2. Crystal structures of the active, open state of SHP2^E76K^ showed an alternative relative arrangement of N-SH2 and PTP that exposes the active site of PTP to the solvent (Figure 1b).^44^ According to the SHP2^E76K^ crystal structure, SHP2 would undergo a huge conformational transition for passing from the inactive to the active state, involving a 120° rotation of the C-SH2 domain and the relocation of the N-SH2 domain to a PTP surface opposite the active site.^44^ However, it is questionable whether this open structure is representative of active SHP2 in solution because the arrangement of the N-SH2 and C-SH2 domains is incompatible with the simultaneous binding of a BTAM with two phosphorylated motifs connected by a linker of 40 Å.^42^ In addition, MD simulations suggested that the open structure of SHP2^E76K^ resolved by crystallography exhibits structural variability in solution,^45^ indicating that further rearrangements may take place.

Despite the utmost role of overly active SHP2 in disease, and despite long-term efforts, the solution ensemble of active SHP2 remains a conundrum. A valuable characterization of the open state of SHP2 has been given by small-angle X-ray scattering (SAXS) experiments. Several SAXS curves of SHP2 are available in literature, and they have provided information on the radii of gyration (*R*_g_) of SHP2 in different conditions.^18, 44, 46^ The *R*_g_ of the constitutively active SHP2^E76K^ mutant in solution is 2.94 nm, a value that is significantly larger than the *R*_g_ of wild-type SHP2 in solution, 2.69 nm.^44, 46^ Therefore, SHP2 in autoinhibited state adopts a more compact conformation than SHP2 in the open state. However, the lack of structural information at atomistic level had so far precluded the characterization of the three-dimensional ensemble of the open state.

In spite of challenges faced by experiments, some key hallmarks of the open, active state of SHP2 are well known. In fact, while the autoinhibited structure of SHP2 is characterized by a tight binding of N-SH2 to PTP and by perfect complementarity between these two domains, the open structure seems to be characterized, instead, by non-specific interactions between N-SH2 and PTP. If an alternative interdomain interface between N-SH2 and PTP were present in the open structure, such interface would represent another site for the modulation of the activation mechanism. However, no mutation has been identified so far in support of an additional interface.^6, 8^ Furthermore, the peaks in the [^1^H-^15^N]-TROSY-HSQC spectrum of the truncated form ΔN-SH2, lacking the N-SH2 domain, were nearly superimposable with the corresponding full-length SHP2^E76K^ peaks, suggesting that the ΔN-SH2 construct perfectly mimics the C-SH2–PTP moiety of the full-length, open SHP2^E76K^.^46^ Finally, the crystal structure of the truncated form ΔN-SH2 showed an approximate 120° rotation of the C-SH2 domain compared to the autoinhibited structure,^46^ exactly like in the crystal structure of open SHP2^E76K^.^44^ All these data indicated that the rearrangement of the C-SH2 domain relative to PTP may occur independently of the presence of N-SH2. Hence, for the modelling of the open SHP2, atomic clashes between N-SH2 and PTP should obviously be avoided, but the unspecific interaction between N-SH2 and PTP might in first instance be neglected.

In this work, we validate structures and conformational ensembles of SHP2 against SAXS data. Since our SAXS predictions take an explicit representation of the hydration layer and excluded solvent,^47, 48^ they do not require fitting of the hydration layer or excluded solvent densities against the experimental data. Furthermore, the effect of the hydration layer on the radius of gyration *R*_g_ detected by SAXS is accurately captured by explicit-solvent MD simulations.^49^ Consequently, small discrepancies of the *R*_g_ between model and experiments in the order of 0.5 Å or even less are detected and not absorbed by fitting parameters.^50^

By means of molecular dynamics (MD) simulations, we have derived a heterogeneous solution ensemble of active SHP2^E76K^. Our results demonstrate that the MD-derived ensemble is in excellent agreement with the SAXS data of SHP2^E76K^ reported by Pádua *et al*.,^46^ obviating the need of any further refinement of the ensemble. The relative arrangement of the tandem SH2 domains (N-SH2–C-SH2) in the proposed SHP2^E76K^ ensemble has been guided by a previously derived tandem SH2 ensemble (*i.e.*, in absence of the PTP domain), validated against SAXS and NMR residual dipolar couplings of N-SH2–C-SH2.^51^ Consequently, the relative N-SH2–C-SH2 arrangement in the SHP2^E76K^ ensemble would allow the binding stimulating bidentate peptides, providing a valuable insight into its functional relevance. Moreover, our strategy to derive the SHP2^E76K^ ensemble is in line with the [^1^H-^15^N]-TROSY-HSQC spectrum of ΔN-SH2 mentioned above. Therefore, the derived ensemble is compatible with available SAXS and NMR data of SHP2^E76K^, tandem SH2, and ΔN-SH2, and it would allow the binding of bidentate peptides, in contrast to the crystal structure of open SHP2^E76K^. The fact that neither the conformational ensemble nor the hydration layer or excluded solvent were fitted against SAXS data excludes any problems owing to overfitting, enhancing the faith in the credibility of the proposed SHP2^E76K^ ensemble.

## RESULTS

### Analysis of SAXS curves from crystal structures of closed SHP2 and open SHP2^E76K^

In this section the experimental SAXS curves of wild-type SHP2 and of the SHP2^E76K^ mutant from Ref. 46 (Figure 1c-d, Figure S1, black dots) are compared with the SAXS curves predicted from MD simulations (Figure 1c-d, Figure S1, red curves) generated starting from the corresponding crystallographic structures^15, 44^ (respectively PDB ID 4DGP and 6CRF, cf. Figure 1a-b). For the sake of completeness, the results of the fitting from coarse-grained rigid-body modelling by Pádua *et al.* are also reported (Figure 1c-d, green curves).^46^ These results provide an overview on the state-of-the-art of the structural interpretation of the scattering data of SHP2 with a single-structure model fitting.

The SAXS curve of wild-type SHP2 agrees reasonably well with the closed crystal structure of autoinhibited SHP2 over the entire *q*-range (Figure 1c). The SAXS curve calculated from explicit-solvent MD simulations (Figure 1c, red curve) returned a radius of gyration of 2.71 nm, in excellent agreement with the experimental value of 2.69 nm. The model fitting by Pádua *et al.* (Figure 1c, green curve) reproduced the experimental curve with similar accuracy.^46^ Therefore, the size and the shape of wild-type SHP2 in closed, autoinhibited conformation, resolved by X-ray diffraction, well match with the SAXS curve of wild-type SHP2 in solution, since the *R*_g_ and many features of the SAXS curve were reproduced. This confirms that, at basal conditions, wild-type SHP2 mostly adopts a closed and compact conformation in solution.^6, 8^

On the other hand, neither the crystal structure of SHP2^E76K^ in open conformation^44^ nor the rigid-body model fitting^46^ has accurately reproduced all features of the SAXS curve of SHP2^E76K^ in solution (Figure 1d). The SAXS curve calculated from explicit-solvent MD simulations (Figure 1d, red curve) restrained to backbone positions of the crystal structure of open SHP2^E76K^ (PDB ID 6CRF^44^) yields a radius of gyration of 2.83 nm, significantly smaller than the experimental value of 2.94 nm. However, the shape of the curve at higher *q*-values was reasonably well reproduced. Instead, the SAXS curve obtained via rigid-body modelling (Figure 1d, green curve)^46^ deviates from the experimental curve at higher *q*-values, where a minimum was reported in place of a smoother decrease of the scattering signal. In exchange, the rigid-body modelling perfectly reproduced the experimental radius of gyration and the shape of the SAXS curve at lower *q*-values,^46^ as expected from a fitted structural model. Furthermore, the model proposed by Pádua *et al.*^46^ differed from the crystal structure resolved by LaRochelle *et al.*^44^ While in the latter the N-SH2 domain is in contact with PTP, in the former N-SH2 is detached from PTP. The ambiguity of these results and the challenges to match the experimental SAXS curve with a single-structure model over the entire *q*-range suggests that the constitutively active SHP2^E76K^ mutant might adopt a range of conformations, *i.e.*, a heterogenous solution ensemble.

### Conformational ensembles of tandem SH2 (N-SH2–C-SH2)

In order to overcome the limitations of the single-structure model fitting, we used MD simulations for generating the solution ensemble of SHP2^E76K^. However, exploring the whole conformational space of active SHP2 is demanding in terms of computational costs. Hence, instead of starting simulations of full-length SHP2^E76K^ (residues Met^1^–Ile^529^) at the crystal structure conformation, we first generated a pool of structures of relative arrangements of *i)* the tandem SH2 domains N-SH2– C-SH2 (SHP2^1-220^, residues Met^1^–Arg^220^),^51^ and *ii)* of truncated SHP2, i.e., C-SH2–PTP lacking the N-SH2 domain (ΔN-SH2, SHP2^105-525^, residues Ala^105^–Leu^525^). This strategy is supported by NMR data showing that interactions between N-SH2 and PTP are unspecific in the open state,^46^ while SHP2^E76K^ almost completely populates the open state.

During the simulations, the tandem SH2 populates a plethora of conformations, with the SH2 domains adopting different relative orientations. A detailed analysis of the tandem SH2 rearrangement during these simulations has been presented elsewhere.^51^ In the previous work, the solution ensemble of the tandem SH2 was successfully compared with the available experimental data from SAXS and NMR residual dipolar couplings.^51^ This allowed us to validate the simulation protocol used here for the truncated and full-length SHP2. In the present work we exclusively focus on the conformations of the tandem SH2 necessary to generate the solution ensemble of SHP2^E76K^.

The flexibility of the linker allows the rearrangement of the tandem SH2, which is mainly accomplished either by the rotation of the SH2 domains around the principal axis of the tandem SH2 or by the bending of the flexible linker, as illustrated by superimposing the C-SH2 domain (Figure 2a). A principal component analysis (PCA) revealed that the interdomain relative fluctuations largely prevail over other internal motions of the domains, since the first five principal components encompassed almost 90% of the overall fluctuations. For the sake of simplicity, we represented the relative fluctuations using a minimal set of two modes of motion (respectively represented by the principal components PC1 and PC2), which were representative of nearly 60% of the overall tandem SH2 fluctuations. Figure 2b presents the free energy landscape of tandem SH2 projected onto the two principal components PC1 and PC2, together with the central structures for the twenty most populated clusters.

**Figure 2.**
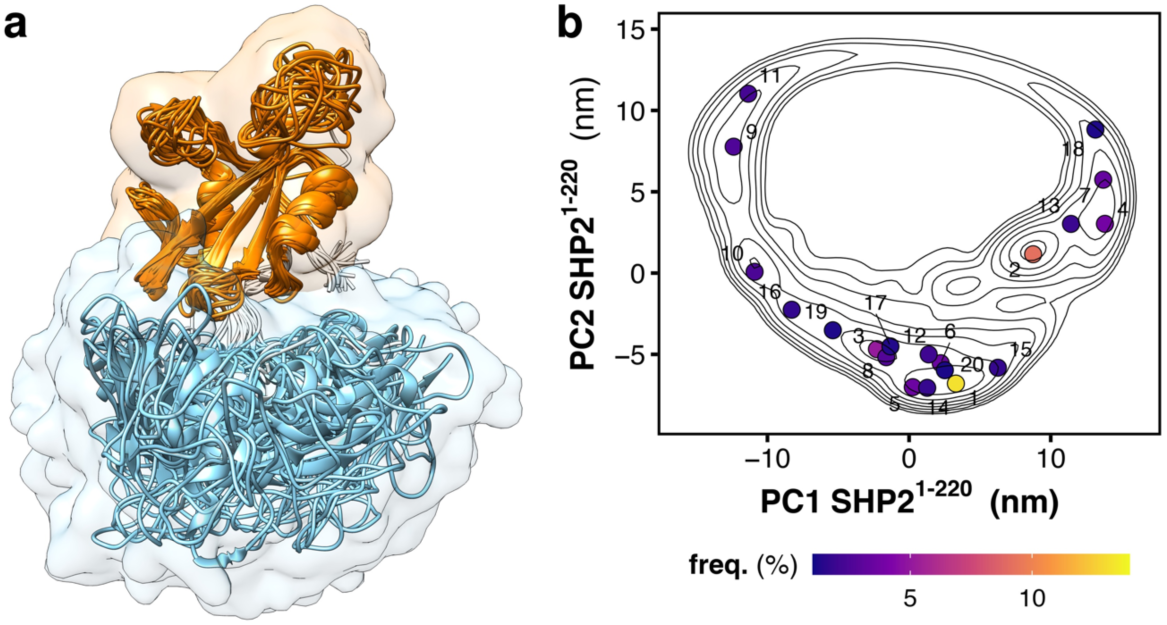
Structural dynamics and free energy landscape of tandem SH2 domains. a) Overlay of the representative structures of the tandem SH2 obtained from MD simulations, as taken from Ref. 51. N-SH2 and C-SH2 are depicted as ribbons and colored respectively in cyan and orange. The structures were superimposed at the C-SH2 domain. b) Free energy landscape along the principal components PC1 and PC2 of the tandem SH2 (SHP2^1-220^) as obtained from MD simulations. The projection of the central structure of the twenty most populated clusters are reported as dots in the essential plane.

### Conformational ensemble of truncated SHP2 (ΔN-SH2)

During the simulations of the truncated SHP2 form, ΔN-SH2, the C-SH2 domain accomplished a roto-translation relative to PTP. This motion is coupled with the torsion and the bending of the linker connecting the C-SH2 domain to PTP. Figure 3 shows three representative structures of ΔN-SH2. These structures correspond either to the conformation adopted by the C-SH2−PTP moiety in autoinhibited SHP2 (Figure 3a), or to the most populated conformations adopted by ΔN-SH2 during the simulations in solution (Figure 3b-c). To reveal the large-scale motions of the C-SH2 domain relative to PTP, a PCA was performed on the rigid core of the C-SH2 domain after least-square fitting over the core of the PTP domain. The first two principal components, PC1 and PC2, encompass 82% of the whole motion of the C-SH2 core. Figure 3d presents the free energy landscape of ΔN-SH2 projected on the essential plane spanned by two principal components PC1 and PC2. Two major basins are visible, separated by a small free energy barrier of less than 5 kJ/mol. The free energy landscape is compatible with the projections of the central structures for the twenty most populated clusters (Figure 3e). A minimum free-energy pathway connecting the two minima (Figure 3d, dotted line) corresponds to the roto-translation of the C-SH2 domain relative to PTP, observed during the simulations (Figure 3a-c, Movie S1).

**Figure 3.**
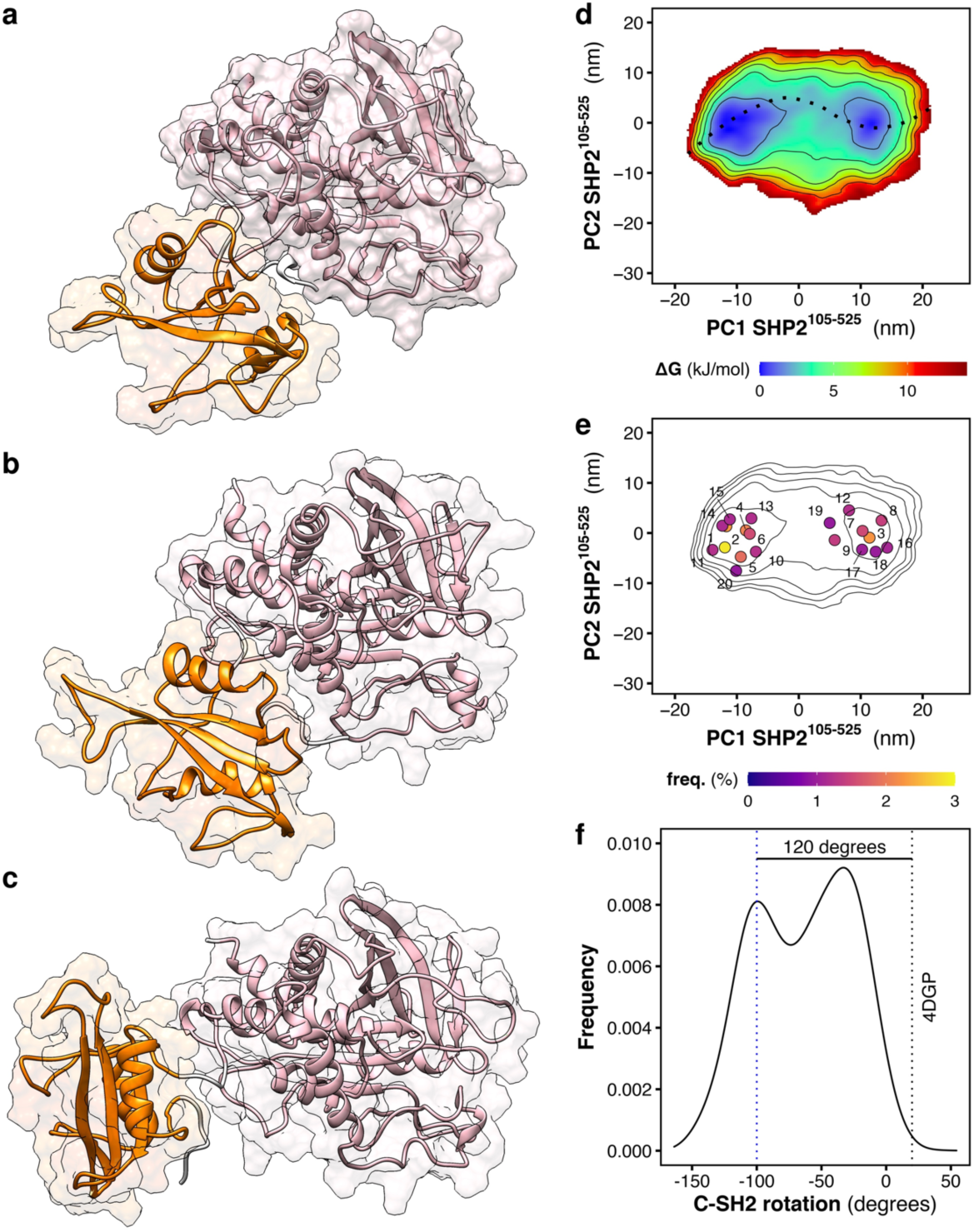
Structural dynamics and free energy landscape of truncated SHP2. Simulation snapshots of truncated SHP2 ΔN-SH2 (SHP2^105-525^): a) conformation adopted by the C-SH2−PTP moiety in autoinhibited SHP2,^15^ b-c) the two most populated conformations adopted by ΔN-SH2 during the simulations. C-SH2 and PTP are depicted as ribbons and colored respectively in orange and pink. d) Free energy landscape along the principal components PC1 and PC2 of truncated SHP2 ΔN-SH2 as obtained from MD simulations. The free-energy minimum pathway, reported over the essential plane as a dashed line, corresponds to the roto-translation of the C-SH2 domain relative to PTP (see also Movie S1). e) Free energy landscape as contour plot together with the central structures of the twenty most populated clusters. f) Distribution of the dihedral angle representing the rotation of the C-SH2 domain relative to PTP. The value of the dihedral angle in autoinhibited SHP2 (PDB ID 4DGP^15^) is reported as a dashed vertical black line. The blue line indicates the value of the dihedral angle that corresponds to the structure depicted in panel c.

Analysis of the hydrogen bonds did not reveal any stable interactions between residues of the C-SH2 domain with PTP throughout the rotation. Instead, a large number of weak interaction pairs are accessible while the frequency of each H-bond is negligible or below 10%. In contrast, rather stable H-bonds (30-60% of frequency) are formed between the residues of the linker connecting C-SH2 to PTP: the bent conformation adopted by the linker after the C-SH2 roto-translation is stabilized by the interaction of the side chain of Asn^217^ with the backbone of residues Thr^219^ and Arg^220^. In addition, the analysis of the Coulomb potential surfaces in the structure with rotated C-SH2 shows a weak long-range interaction of the negatively charged patch on the PTP surface, formed by Glu^225^, Glu^227^, and Asp^485^, with the positively charged patch of the C-SH2 surface, formed by Lys^131^ and Lys^198^. In conclusion, the C-SH2 displacement does not result in a network of stable interactions at the interface between C-SH2 and PTP, and the roto-translation of C-SH2 involves a transition over a low free energy barrier.

Apart the essential eigenvectors, the rotation of the C-SH2 domain relative to PTP was monitored by means of the dihedral angle defined by the positions of the Cα atoms of residues Glu^139^−Ser^134^−Pro^454^−Cys^459^ (Figure S2). These residues represent the ends of the βB strand of C-SH2 and of the βM strand of PTP, which resides approximately at the center of the respective domains and adopt an orientation roughly perpendicular to the axis connecting the center-of-mass of the two domains. The distribution of the dihedral angle (Figure 3f) is bimodal and the two peaks respectively correspond to the basins reported in the essential plane (Figure 3d). The value of the dihedral angle is reported also for the configuration of the C-SH2−PTP moiety in autoinhibited structure of SHP2 (Figure 3f, dotted black line, 20 degrees). Critically, such a conformation is hardly explored by ΔN-SH2 in solution. One of the peaks is placed at −100 degrees (Figure 3f, dotted blue line), just 120 degrees from the orientation of C-SH2 in autoinhibited SHP2, in agreement with the observations by NMR.^46^ The second peak at −30 degrees corresponds to an intermediate conformation not previously detected by experiments (Figure 3b).

### Heterogeneous ensemble of full-length SHP2

As described above, MD simulations revealed the configurational space of tandem SH2^51^ and of ΔΝ-SH2. Next, we combined these fragments to obtain a large set of initial conformations of open SHP2^E76K^. The first 20 clusters respectively encompass 64% of the conformations of the tandem SH2 and 30% of the conformations of the ΔN-SH2 construct. In addition, the clusters are distributed over the whole essential plane, therefore they are representative of all principal conformational states of both tandem SH2 and ΔN-SH2. All 20 times 20 pair-wise combinations of the representative structures of tandem SH2 and of the ΔN-SH2 construct were analyzed with respect to clashes between PTP and N-SH2 (Figure 4). After the superposition of each representative structure pair at the C-SH2 domain, the number of clashes raising between the N-SH2 and the PTP domain was determined. If the number of clashes did not exceed 100, the representative structure pair was further considered as a template for the following modelling of the full-length SHP2^E76K^ ensemble (Figure 4a). The comparative protein structure modelling, which generated the full-length SHP2^E76K^ conformations, was based on the satisfaction of spatial restraints. Indeed, the relative positions of the domains in the template structures affected the final model. Among the 400 possible pairs, 181 representative structure pairs had no clashes, 275 pairs had fewer than 100 clashes, while 125 pairs were discarded since they had at least 100 clashes (Figure 4b). The threshold of 100 for the number of clashes was arbitrarily chosen, yet considering that such a number was an indication of only a moderate overlap between the domains, mainly caused by surface residue side chains, and that such contacts between the domains are readily corrected during the modelling phase.

**Figure 4.**
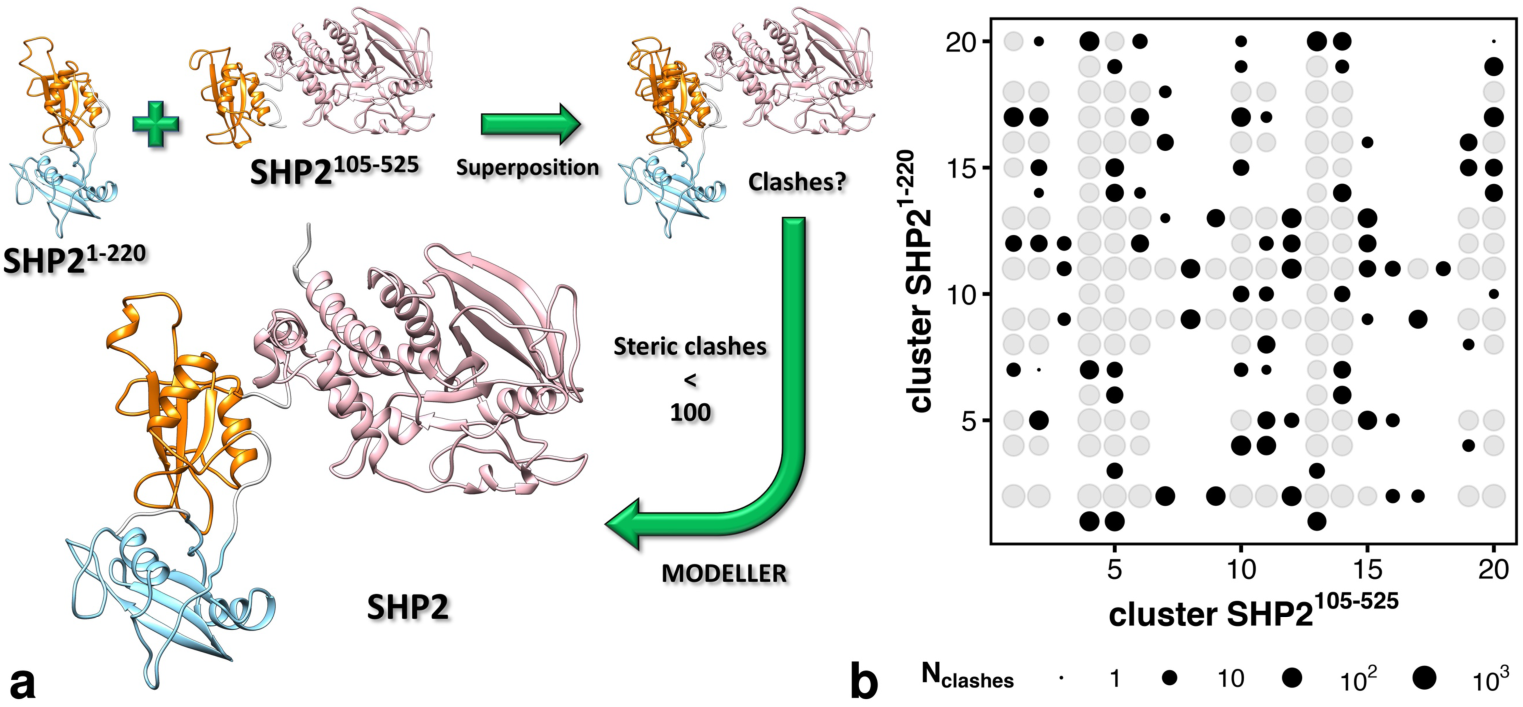
Homology modeling workflow for generating SHP2^E76K^ conformations. a) Scheme representing the homology modelling flowchart used for the generation of the open conformations of SHP2^E76K^ used as seeds in MD simulations. Representative structures of the tandem SH2 (SHP2^1-220^) and of truncated SHP2 ΔN-SH2 (SHP2^105-525^) were superposed at the C-SH2 domain. After the superposition, the number of clashes raising between PTP and N-SH2 was determined. If the number of clashes did not exceed 100, the representative structure pair was further used as a template for the following modelling of the full-length SHP2^E76K^ (SHP2^1-529^). b) Number of clashes between PTP and N-SH2 for each pair of representative structures of the tandem SH2 (SHP2^1-220^) and of truncated SHP2 ΔN-SH2 (SHP2^105-525^). Among the 400 possible pairs, 181 representative structure pairs had no clashes (void spaces), 275 pairs had fewer than 100 clashes (void spaces and black dots), 125 pairs had at least 100 clashes (gray dots) and were discarded from homology modelling.

The 275 models of full-length SHP2^E76K^ generated by homology displayed a high structural variability (Movie S2) with the radius of gyration *R*_g_ spanning from 2.6 nm to 3.4 nm (Figure S3). To facilitate the comparison of *i) R*_g_ of the bare individual structures generated by homology with *ii)* the *R*_g_ of the ensemble obtained via explicit-solvent SAXS calculations, we increased the *R*_g_ of individual models by a fixed value of 0.76 Å, corresponding to the increase of *R*_g_ owing to the hydration layer (*vide infra*).

The 275 models of full-length SHP2^E76K^ generated by homology were used as seeds for 100 ns MD simulations per model. After the equilibration, the conformations sampled in the last 10 ns chunk of the simulations were collected and aggregated into a cumulative trajectory (2.75 μs) representing the solution ensemble of open SHP2^E76K^.

According to the derived solution ensemble, SHP2^E76K^ adopts a plethora of conformations. The flexibility of the linkers connecting N-SH2 to the C-SH2 domain, and connecting C-SH2 to the PTP domain, allow for large-scale conformational rearrangements, which include the wide relocation of the N-SH2 domain relative to PTP, as illustrated by superposing the structures of SHP2^E76K^ at the PTP domain (Figure 5a). Notably, the solution ensemble of SHP2^E76K^ comprises structures where the N-SH2 domain is still relatively close to the surface of the PTP domain or nearby the active site, as well as structures where the N-SH2 domain is completely detached from PTP or far apart from the active site (Movie S2).

**Figure 5.**
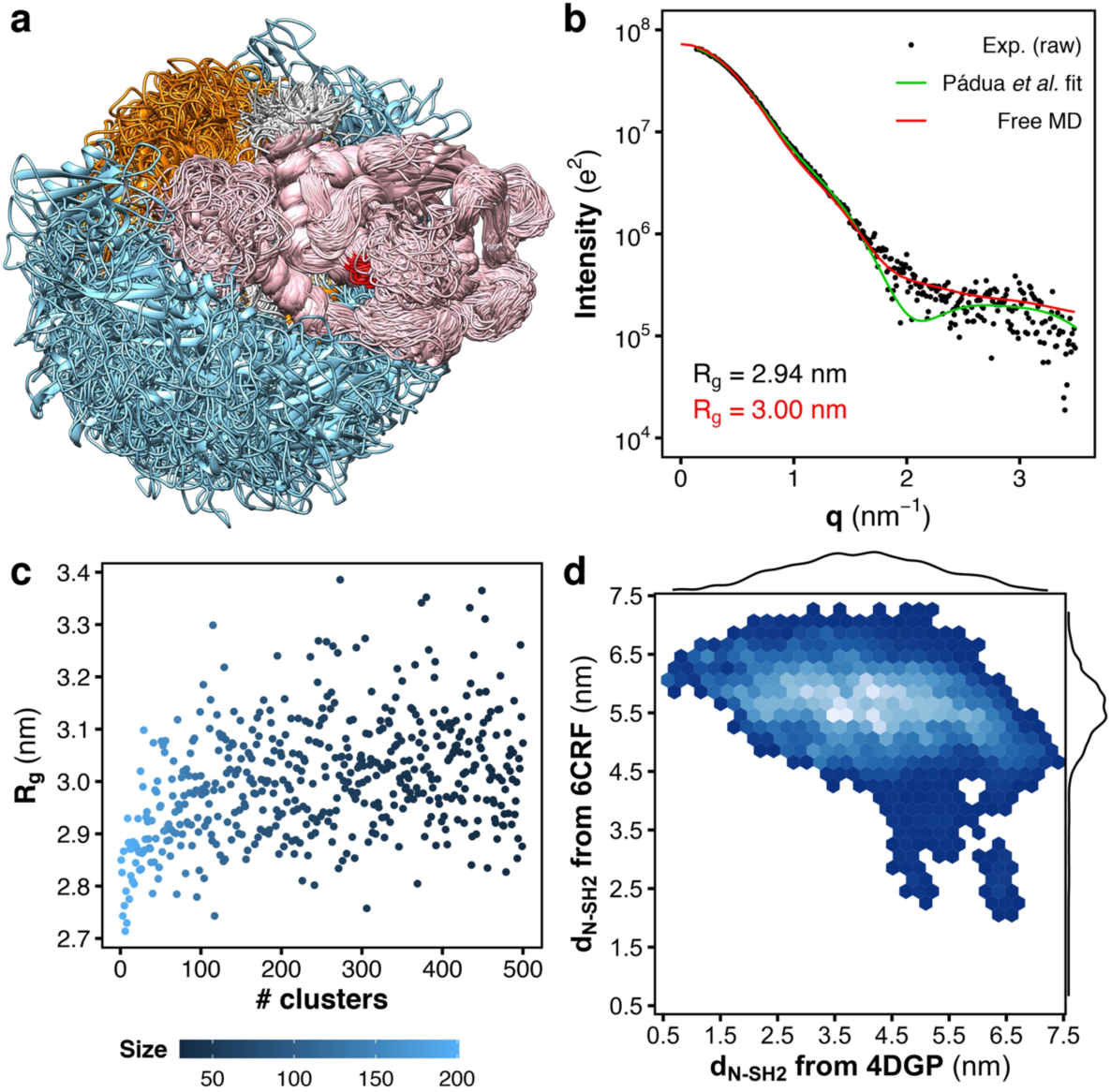
Heterogeneity of the SHP2^E76K^ structural ensemble is supported by MD simulations and SAXS. a) Overlay of representative structures of SHP2^E76K^ as obtained from MD simulations. N-SH2, C-SH2, and PTP are depicted as ribbons and colored respectively in cyan, orange, and pink. The catalytic site is highlighted in red. The structures were superposed at the PTP domain. b) Small-angle X-ray scattering (SAXS) curve calculated from the solution ensemble of SHP2^E76K^ (red line) is compared with the experimental curve (black dots), reported as raw data, and with the single-structure model by Pádua *et al*. (green line).^46^ Radii of gyration (*R*_g_) from the MD ensemble and from experiment are reported with black and red font, respectively. c) *R*_g_ values of SHP2^E76K^ calculated from the ensembles of the first 500 most populated clusters via Guinier fit to calculated SAXS curves. The population of the clusters are reported in color scale. d) Density map of the distance of the N-SH2 center-of-mass (*d*_N-SH2_) from the positions respectively occupied by the N-SH2 center-of-mass either in the crystal structure of autoinhibited SHP2 (4DGP^15^) or in the crystal structure of open SHP2^E76K^ (6CRF^44^). The distributions of the distance *d*_N-SH2_ from 4DGP and from 6CRF are reported as marginal plots.

In order to validate the solution ensemble obtained from MD simulations, the SAXS curve was calculated using accurate explicit-solvent SAXS curve predictions.^47, 48^ The calculated SAXS curve was compared both with the experimental raw data (Figure 5b, Figure S4, black dots) and with the SAXS curve obtained from the single-structure model by Pádua *et al*. (Figure 5b. green curve).^46^ The calculated SAXS curve obtained from the MD solution ensemble (Figure 5b, red curve) is in excellent agreement with the experimental data. The calculated SAXS curve overestimated the radius of gyration only marginally by 0.6 Å. Such agreement is remarkable considering that neither the structural ensemble nor the hydration layer or the excluded volume were refined or fitted against the SAXS data, providing high confidence in the validity of the ensemble. In addition, the calculated SAXS curve well reproduced the scattering signal decay of the experimental SAXS curve at higher *q*-values, on the contrary of SAXS curve from the single-structure model by Pádua *et al.*,^46^ which instead reported a minimum at q ≈ 2 nm^−1^ (Figure 5b, green curve). Notably, the presence of minima at higher *q*-values is an indication of a protein with a distinct shape or of a more homogeneous ensemble. The lack of such minima in the experimental data indicates that SHP2^E76K^ does not adopt a single structure in solution but instead a heterogenous ensemble, in agreement with the MD-derived structural ensemble.

To assess the impact of local relaxation on the description of the SAXS curve and validate the robustness of the final ensemble, we calculated SAXS curves using explicit-solvent SAXS curve predictions applied to the cumulative trajectories of other 10 ns intervals of the simulations. The SAXS curve from the first 10 ns of the simulations (Figure S5, red curve) closely matched the curve obtained from the last 10 ns of the simulations (Figure 5b, red curve). Similar convergence was observed in other cumulative trajectories (Figure S6), producing SAXS curves with an average radius of gyration (*R*_g_) of ∼3.00 nm and an average increase of *R*_g_ owing to the hydration layer by 0.76 Å. These findings demonstrate the excellent convergence of the simulations in describing the SAXS curve of SHP2^E76K^.

To quantify the heterogeneity of the SHP2^E76K^ ensemble, we derived the cumulative frequency of the SHP2^E76K^ conformations as a function of the number of clusters. In fact, the first 20, 100, 200, 300, 400, and 500 most populated clusters encompass respectively 7%, 31%, 50%, 62%, 70%, and 77% of total conformations, characterizing a highly heterogeneous ensemble (Figure S7). Furthermore, we determined the range of radii of gyration (*R*_g_) adopted by SHP2^E76K^ in solution by computing SAXS curves from the ensembles of the first 500 most populated clusters (Figure 5c, Movie S3). The *R*_g_ values were obtained by Guinier fits to calculated SAXS curves, thus the values include contributions from the hydration layer. The *R*_g_ values of the clusters demonstrate that SHP2^E76K^ may adopt compact conformations in solution, with a *R*_g_ of only 2.7 nm, close to the *R*_g_ observed in the autoinhibited structure of wild-type SHP2. However, in these compact solution structures the active site is exposed to the solvent, yet the N-SH2 domain is in contact with the PTP domain. The largest *R*_g_ adopted by SHP2^E76K^ is nearly 3.4 nm, corresponding to wide open and elongated structures with the N-SH2 domain located at large distance from PTP. Hence, the *R*_g_ of ∼3.0 nm observed in the SAXS curve in Figure 5b is an ensemble-averaged value that does not reflect the spatial extension of all possible structures of the heterogeneous ensemble.

To further characterize the solution ensemble of SHP2^E76K^, we considered the distance of the N-SH2 center-of-mass (*d*_N-SH2_) from the positions respectively occupied by the N-SH2 center-of-mass either in the crystal structure of autoinhibited SHP2 (4DGP) or in the crystal structure of open SHP2^E76K^ (6CRF). The distance *d*_N-SH2_ from the 4DGP position continuously ranges from 0.5 to 7.4 nm, with a modal value of ∼4.0 nm (Figure 5d), confirming that the solution ensemble includes configurations of SHP2^E76K^ with a diverse degree of opening. The distance *d*_N-SH2_ from the 6CRF position ranges from 2.0 to 7.3 nm, with a modal value of ∼5.5 nm, indicating that the N-SH2 domain had never reached the position occupied in the crystal structure of open SHP2^E76K^ (6CRF), though it is close to it with low probability.

The three-dimensional density distribution of the center-of-mass position of the N-SH2 domain relative to PTP (Figure 6a) shows that the position opposite to the active site, occupied by the N-SH2 domain in the crystal structure of SHP2^E76K^ after its relocation, is only one of the many possible positions occupied by N-SH2 in the solution ensemble. We calculated the root mean squared deviation (RMSD) from the respective crystal structures, 4DGP and 6CRF (Figure S8). Critically, most of the conformations of the solution ensemble largely differ both from the autoinhibited structure of SHP2 and from the crystal structure of open SHP2^E76K^. Figure 6b compares the crystal structure of open SHP2^E76K^ with the most similar MD configuration. Although the tandem SH2 in these two structures adopts quite different conformations, the centroids of the domains respectively occupy nearly the same regions of the space.

**Figure 6.**
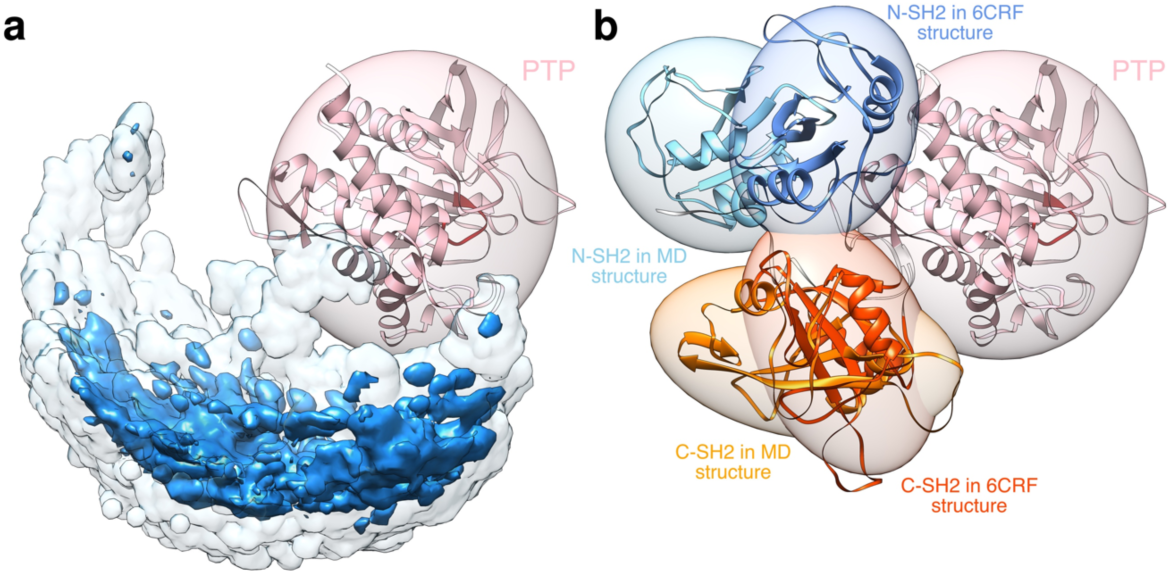
Comparison of the N-SH2 position in MD simulations and crystal structure. a) Three-dimensional density distribution of the center-of-mass position of the N-SH2 domain relative to PTP. The isovalues of the opaque dark cyan isosurface and of the transparent light cyan isosurface are respectively one tenth and one thousandth of the maximum density value. b) Comparison of the crystal structure of open SHP2^E76K^ with the most similar MD configuration. N-SH2, C-SH2, and PTP are depicted as ribbons and colored respectively in cyan, orange, and pink (see text labels). The catalytic site is highlighted in red. The inertia ellipsoid of each domain is reported as a transparent surface. The structures were superposed at the PTP domain.

## CONCLUSIONS

We combined MD simulations, homology modelling, and explicit-solvent SAXS curve predictions to derive and validate the heterogeneous atomistic ensemble of constitutively active SHP2^E76K^ in solution. The protein adopted a plethora of conformations, with radii of gyration ranging between 2.7 and 3.4 nm. Some conformations with N-SH2 still in contact with PTP endowed a radius of gyration close to the value observed for the autoinhibited structure of wild-type SHP2, although the active site was visibly exposed to the solvent. Other structures, instead, were wide open and elongated with the N-SH2 domain far apart from PTP. Notably, the radius of gyration observed in the experimental SAXS curve was an averaged-ensemble value that does not reflect the size of all possible structures of the heterogeneous ensemble. Based on the structural ensemble of SHP2^E76K^ presented here, we anticipate that other constitutively active pathogenic SHP2 mutants may likewise adopt heterogenous ensembles in solution.

The ensemble proposed here is compatible with several lines of experimental data. First, the SHP2^E76K^ ensemble has been successfully validated against experimental SAXS data of SHP2^E76K^.^46^ Second, the initial N-SH2–C-SH2 arrangements for MD simulations were taken from an ensemble of the tandem SH2 systems (in absence of PTP), which was validated previously against SAXS data and NMR residual dipolar couplings.^51^ These relative N-SH2– C-SH2 arrangements are in principle compatible with simultaneous binding of a BTAM with two phosphorylated motifs connected by a linker of 40 Å, in contrast with the N-SH2–C-SH2 arrangement in the crystal structure of open SHP2^E76K^.^44, 51^ Third, we constructed the SHP2^E76K^ ensemble by merging the conformations of two SHP2 fragments, N-SH2–C-SH2 and ΔN-SH2, thus initially neglecting putative interactions between N-SH2 and PTP; this methodology aligns with the absence of specific contacts between N-SH2 and PTP as suggested by comparing the NMR signals of SHP2^E76K^ with those of ΔN-SH2.^46^ Consequently, our heterogeneous ensemble is compatible with available SAXS and NMR data for SHP2^E76K^, tandem SH2, and ΔN-SH2, and the ensemble enables the simultaneous binding of a BTAM to both N-SH2 an C-SH2.

In conclusion, our findings represent a major advancement in our comprehension of SHP2 conformational landscape. Instead of aiming to describe active SHP2 as a single most representative structure, we here describe active SHP2 as a dynamic heterogeneous ensemble. We anticipate that the ensemble-based perspective will be essential for understanding SHP2 signaling and the regulation of SHP2 in response to therapeutic agents.

## METHODS

### MD simulation parameters

All MD simulations were performed with the GROMACS software package (version 2020),^52^ using AMBER99SBws force field.^53, 54^ Long range electrostatic interactions were calculated with the particle-mesh Ewald (PME) approach.^55^ A cutoff of 1.2 nm was applied to the direct-space Coulomb and Lennard-Jones interactions. Bond lengths and angles of water molecules were constrained with the SETTLE algorithm,^56^ and all other bonds were constrained with LINCS.^57^ In the productive runs, the pressure was set to 1 bar using the Parrinello-Rahman barostat,^58^ while the temperature was controlled at 298 K using velocity rescaling with a stochastic term.^59^

The MD ensemble of tandem SH2 was taken from our recent study.^51^ The PCA was performed on the Cα atoms of residues Thr^12^−Pro^215^ (hence excluding the flexible termini).

### MD simulations of truncated SHP2 in solution

Molecular dynamics simulations were performed on the truncated SHP2 form lacking the N-SH2 domain (ΔN-SH2) in apo form (SHP2^105-525^, corresponding to sequence ranges from Ala^105^ to Leu^525^). The initial atomic coordinates were derived from crystallographic structure of wild-type SHP2 in autoinhibited conformation (PDB ID 4DGP).^15^ After removing the N-SH2 domain, the residue Ala^105^ was capped. Missing or incomplete residues (strands Leu^236^– Gln^245^, Gly^295^–Val^301^, Phe^314^–Pro^323^; residue Lys^235^) were modeled by Molecular Operative Environment (MOE).^60^ Truncated SHP2 was put at the center of a dodecahedron box, large enough to contain the protein and at least 2.1 nm of solvent on all sides. The system was solvated with ∼49000 explicit TIP4P/2005s water molecules,^61^ and four Na^+^ ions were added to neutralize the simulation box. Then, the solvent was relaxed and the system minimized. After generating the initial velocities at 50 K from different seeds according to the Maxwell distribution, 24 simulations were spawned. The temperature was thermalized to 298 K in 10 ns, in a stepwise manner, while the pressure was set to 1 bar using the weak-coupling barostat.^62^ Finally, 24 independent production simulations of 1.05 μs were performed. The first 50 ns of each simulation and the misfolded structures were discarded, and the analyses were performed on a cumulative trajectory of 23 μs.

The PCA was performed on the rigid core of the C-SH2 domain (Cα atoms of residues Gly^119^−Lys^129^, Ser^134^−Arg^138^, Phe^147^−Arg^152^, Val^167^−Ile^172^, Leu^190^−Lys^199^) after the least-square fitting over the core of the PTP domain (backbone of residues Ser^326^−Cys^333^, Thr^337^−Thr^356^, Tyr^370^−Trp^371^, Lys^378^, Met^383^−Asn^387^, Lys^389^, Ser^391^, Tyr^396^−Ser^404^, Arg^413^−Phe^420^, Pro^432^−Glu^447^, Gly^453^−Ser^460^, Gly^464^−Leu^475^, Asp^489^−Arg^501^, Gln^510^−Tyr^521^).

### MD simulations of open SHP2^E76K^ in solution

Molecular dynamics simulations were performed on SHP2^E76K^ in apo form (SHP2^1-529^, corresponding to sequence ranges from Met^1^ to Ile^529^), excepting for the N-terminus that was modified prepending three residues, Gly^-2^, Ser^-1^ and Gly^0^. Multidomain protein structures of open SHP2^E76K^ were assembled by homology using MODELLER.^63^ The first 20 representative structures from MD simulations of the tandem SH2, ranging from Gly^-1^ to Gln^214^, and of the truncated SHP2 (βN-SH2), ranging from Ser^118^ to Leu^525^, were used as templates. Residues from Leu^525^ to Ile^529^ were modelled in α-helix. All 20×20 combinations between the structures of tandem SH2 and of truncated SHP2 were tried. Combinations that returned more than 100 atom-pair clashes between N-SH2 and PTP were discarded, after the superposition of representative structures at the C-SH2 domain. For each suitable combination, 275 in total, up to 15 independent models were generated, and assessed by the GA341 method,^64^ and by the DOPE (Discrete Optimized Protein Energy) method.^65^ Finally, for each of 275 combinations, the model with the best DOPE score was chosen. Each structure of open SHP2^E76K^ was put at the center of a dodecahedron box, large enough to contain the protein and at least 2.1 nm of solvent on all sides. Each system was solvated with ∼94700 explicit TIP4P/2005s water molecules,^61^ and one Cl^−^ ion was added to neutralize the simulation box. Then, the solvent was relaxed and each system minimized. After generating the initial velocities at 50 K from different seeds according to the Maxwell distribution, the temperature was thermalized to 298 K in 10 ns, in a stepwise manner, while the pressure was set to 1 bar using the weak-coupling barostat.^62^ Finally, 275 independent production simulations of 100 ns were performed. The first 90 ns of each simulation and the misfolded structures were discarded, and the analyses were performed on a cumulative trajectory of 2.65 μs.

The cluster analysis was performed considering all Cα atoms of SHP2^E76K^ after least-square fitting over the core of the PTP domain. Starting from each representative structure of the first 500 most populated clusters, a MD simulation of 11 ns was spawned while restraining some folded regions of the protein (Cα atoms of residues Thr^12^−Gly^24^, Ser^28^−Ser^34^, Phe^41^−Glu^83^, Gly^119^−Lys^129^, Ser^134^−Arg^138^, Phe^147^−Arg^152^, Val^167^−Lys^199^, Trp^248^−Leu^261^, Arg^265^−Arg^278^, Ala^307^−Met^311^, Tyr^327^−Gln^331^, Gln^335^−Asn^349^, Val^352^−Thr^356^, Leu^377^−Arg^421^, Pro^432^−Ile^449^, Val^455^−Cys^459^, Arg^465^−Gln^526^). After discarding the first 1 ns, each trajectory was used for the calculation of the SAXS curve.

### MD simulations of autoinhibited wild-type SHP2 in solution

Molecular dynamics simulations were performed on wild-type SHP2 in apo form (SHP2^1-529^, UniProt Q06124-2, corresponding to sequence ranges from Met^1^ to Ile^529^), except for the N-terminus that was modified prepending three residues, Gly^-2^, Ser^-1^ and Gly^0^. The initial atomic coordinates were derived from crystallographic structure of wild-type SHP2 in autoinhibited conformation (PDB ID 4DGP).^15^ Missing or incomplete residues (strands Gly^−2^– Thr^2^, Leu^236^–Gln^245^, Gly^295^–Val^301^, Phe^314^–Pro^323^, Arg^527^–Ile^529^; residue Lys^235^) were modeled by Molecular Operative Environment (MOE).^60^ The protein was solvated with ∼94700 explicit TIP4P/2005 water molecules,^61^ and one Na^+^ ion was added to neutralize the simulation box. Then, the solvent was relaxed and the system minimized. After generating the initial velocities at 50 K from different seeds according to the Maxwell distribution, 24 simulations were spawned. The temperature was thermalized to 298 K in 10 ns, in a stepwise manner, while the pressure was set to 1 bar using the weak-coupling barostat.^62^ Finally, 24 independent production simulations of 100 ns were performed. The first 90 ns of each simulation were discarded, and the analyses were performed on a cumulative trajectory of 240 ns.

### SAXS calculations from explicit solvent MD simulations

SAXS curves were computed from the MD simulations using explicit-solvent SAXS calculations.^47^ Accordingly, all explicit water molecules and ions within a predefined distance from the protein contributed to the SAXS calculations, as defined by a spatial envelope.^47^ A distance of the envelope from the solute atoms (7 Å) was chosen to ensure nearly bulk-like water at the envelope surface. A density correction was applied to fix the bulk water density to the experimental value of 334 e nm^-3^.^47^ The buffer-subtracted SAXS curve was computed from the scattering of atoms inside the envelope volume, as taken from MD simulation frames of two systems: *i)* containing the protein in solvent and *ii)* containing pure solvent.^47^ The orientational average was carried out using 2000 **q**-vectors per absolute value of *q*. The source code for explicit-solvent SAXS calculations is available at https://gitlab.com/cbjh/gromacs-swaxs. For more details on explicit-solvent SAXS calculations we refer to recent reviews.^50, 66^

The experimental SAXS data for wild-type SHP2 and for SHP2^E76K^, along with the corresponding fits and models, were taken from the Small Angle Scattering Biological Data Bank (SASBDB)^67^ under accession codes SASDEN4 and SASDEP4.^46^

### SAXS calculations from crystal structure of open SHP2^E76K^

Single point SAXS curve calculations were performed by WAXSiS^48^ on SHP2^E76K^ in apo form (SHP2^1-529^, corresponding to sequence ranges from Met^1^ to Ile^529^), excepting for the N-terminus that was modified prepending three residues, Gly^-2^, Ser^-1^ and Gly^0^. The initial atomic coordinates were derived from crystallographic structure of leukemia associated SHP2^E76K^ in open conformation (PDB ID 6CRF, chain A). Missing or incomplete residues (strands Gly^−2^–Ser^−1^, Glu^90^–Asn^92^, Glu^123^–Leu^125^, Thr^127^–Lys^129^, Arg^138^–Asp^146^, Gly^154^–Ser^165^, Lys^166^–Thr^168^, Val^170^–Cys^174^, Gln^175^–Leu^177^, Lys^178^–Val^181^, Glu^185^–Ser^189^, Thr^191^–Asn^200^, Met^202^–Gln^214^, Ala^237^–Lys^244^, Glu^313^–Lys^324^, Gln^526^–Ile^529^; residues Ser^118^, Lys^120^, Lys^131^, His^132^, Ser^134^, Leu^149^, Val^151^, Arg^152^) were modeled by Molecular Operative Environment (MOE).^60^

## AUTHOR CONTRIBUTIONS

M.A. conceived the research. M.A. designed and conducted MD simulations, performed analysis, and compared the results with experimental data. M.A. and J.S.H. designed and supervised SAXS calculations. M.A. drafted the manuscript, and all authors contributed to the editing of the final version.

## DATA AVAILABILITY

Structural models and ensembles generated during the present study are available in the Zenodo repository, DOI 10.5281/zenodo.10057715.

## ACKNOWLEDGEMENTS

This study has received funding from the Deutsche Forschungsgemeinschaft (grant numbers HU 1971/3-1 and INST 256/539-1).

