## Supplementary Information for "Atomistic ensemble of active SHP2 phosphatase"

This PDF file includes:
Figures S1 to S8
References

Other supplementary materials for this manuscript include the following:
Movies S1 to S3

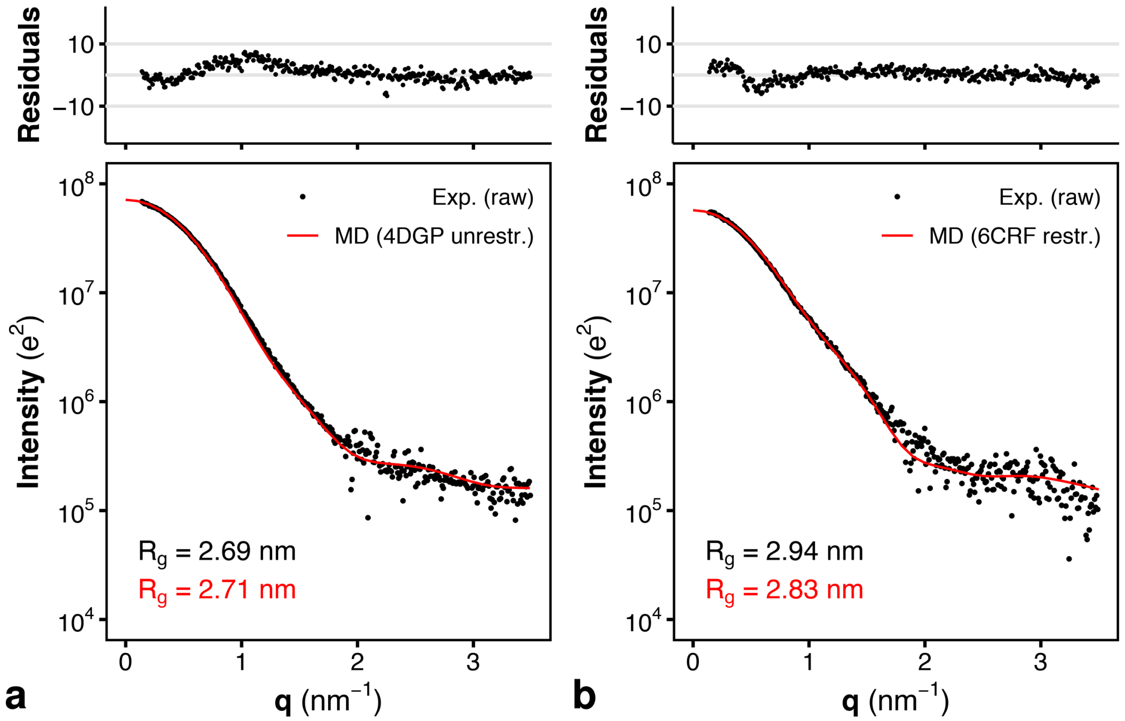


**Figure S1.** Comparison of the small-angle X-ray scattering (SAXS) curve calculated a) from the solution ensemble of the autoinhibited wild-type SHP2 (unrestrained MD simulation from PDB ID 4DGP,[1] red line, $\chi = 3.7$), and b) from the crystal structure of the constitutively active SHP2$^{E76K}$ mutant (restrained MD simulation from PDB ID 6CRF,[3] red line, $\chi = 1.9$), respectively with the corresponding experimental curves (black dots), reported as raw data.[2] Experimental and calculated radii of gyrations ($R_g$) are reported with black and red font, respectively, in each panel. Residuals are plotted as a function of $q$ in the top panels.

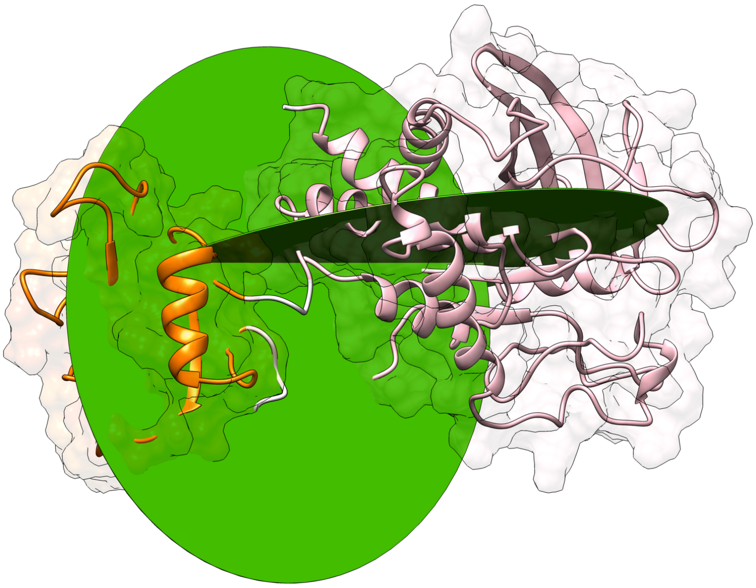

**Figure S2.** Simulation snapshot of truncated SHP2 ΔN-SH2 ($SHP2^{105\text{-}525}$) in one of the most populated conformations adopted by ΔN-SH2 during the simulations. C-SH2 and PTP are depicted as ribbons and colored respectively in orange and pink. The two planes including the dihedral angle defined by the positions of the Cα atoms of residues $Glu^{139}$–$Ser^{134}$–$Pro^{454}$–$Cys^{459}$ are shown in green. These residues represent the ends of the βB strand of C-SH2 and of the βM strand of PTP.

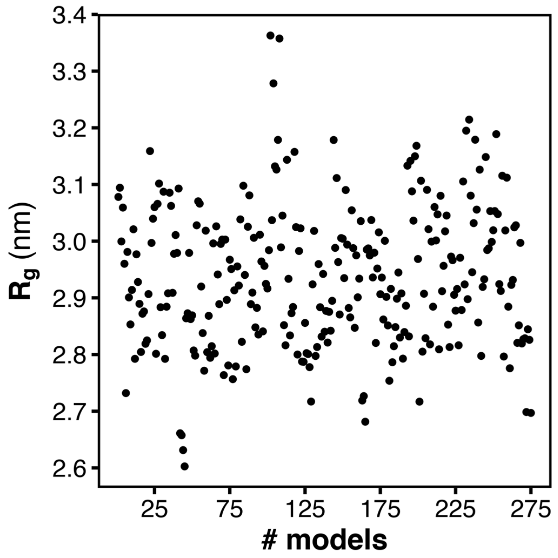


**Figure S3.** Radii of gyration ($R_g$) of the 275 models of full-length SHP2$^{E76K}$ generated by homology modelling. To facilitate the comparison with the $R_g$ values of the structural ensemble obtained via explicit-solvent SAXS calculations, we increased the $R_g$ of individual models by a fixed value of 0.76 Å, corresponding to the thickness of the hydration layer measured from MD simulations.

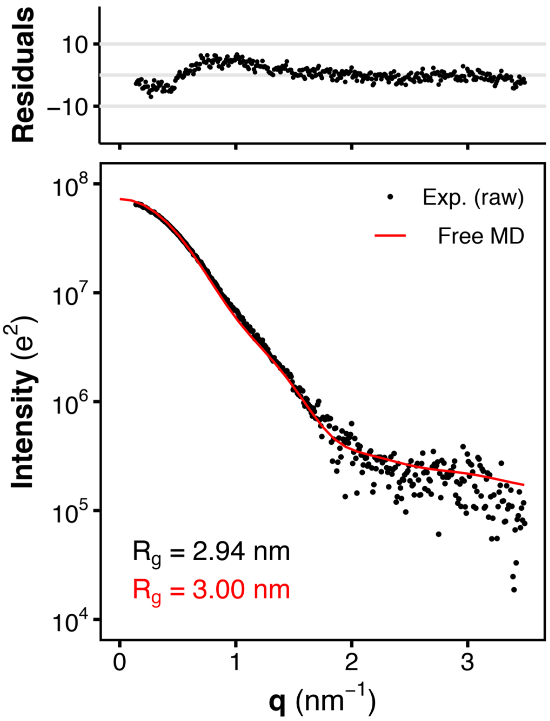


**Figure S4.** Comparison of the small-angle X-ray scattering (SAXS) curve calculated from the solution ensemble of SHP2$^{E76K}$ (red line, $\chi$ = 3.0) with the experimental curve (black dots), reported as raw data, Experimental and calculated radii of gyrations ($R_g$) are reported with black and red font, respectively. Residuals are plotted as a function of $q$ in the top panel.

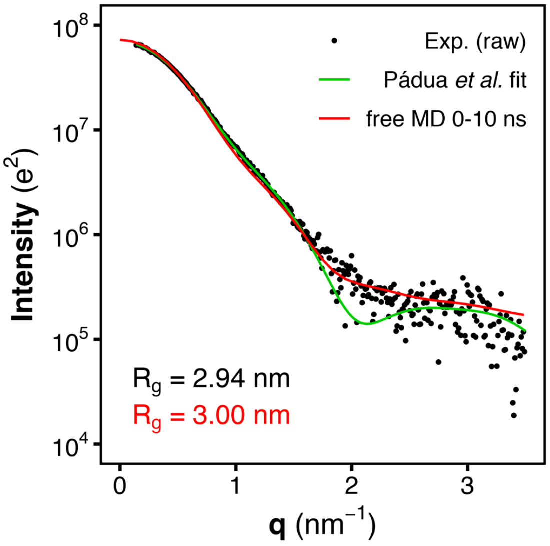


**Figure S5.** Small-angle X-ray scattering (SAXS) curve calculated from the cumulative trajectory of the first 10 ns chunk of simulations of $SHP2^{E76K}$ (red line) is compared with the experimental curve (black dots), reported as raw data, and with the single-structure model by Pádua *et al*. (green line).[2] Radii of gyration ($R_g$) from the MD ensemble and from experiment are reported with black and red font, respectively.

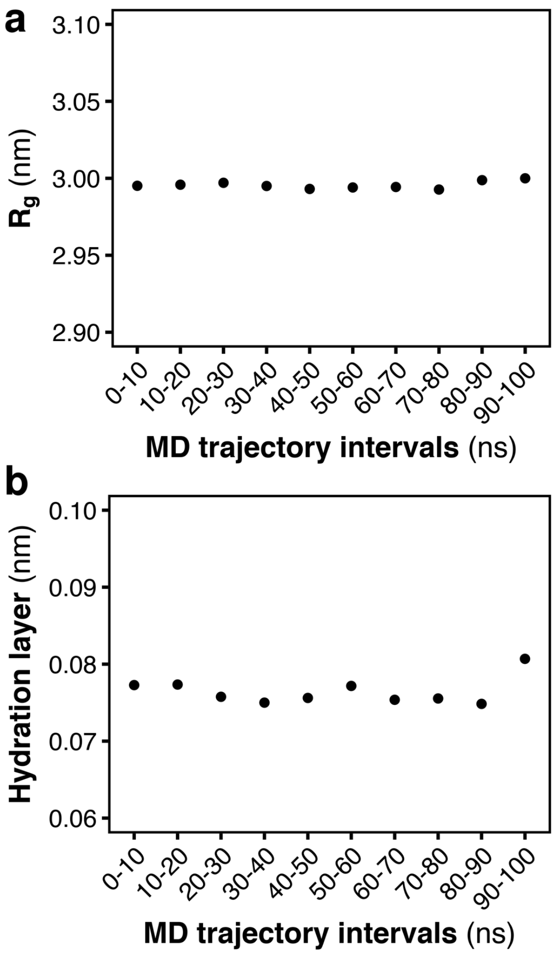


**Figure S6.** Assessment of the impact of local relaxation on the description of the SAXS curve and validation the robustness of the final ensemble: a) radii of gyration ($R_g$), and b) increases of the radius of gyration ($R_g$) owing to the hydration layer, calculated from the cumulative trajectories of SHP2$^{E76K}$ respectively aggregating each 10 ns interval of the simulations.

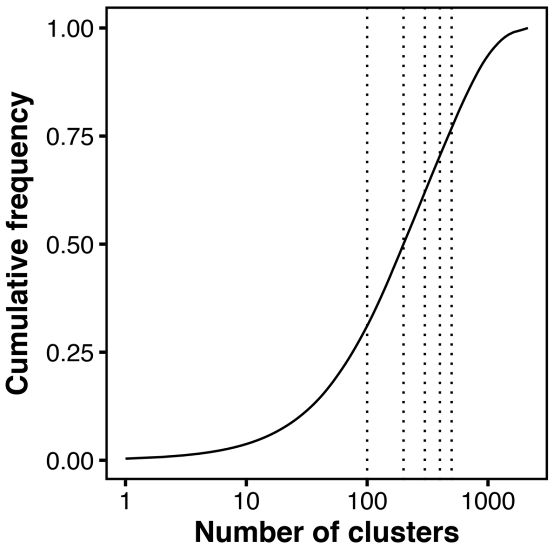


**Figure S7.** Cumulative frequency of cluster population as a function of the number of most populated clusters.

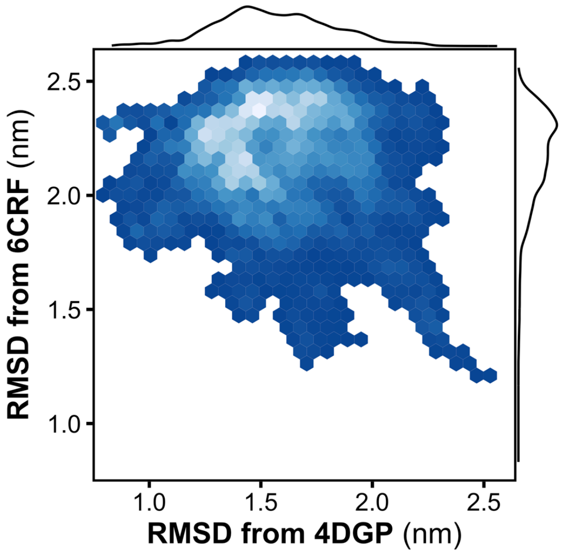


**Figure S8.** Density map of the root mean squared deviation (RMSD) respectively from the crystal structure of autoinhibited SHP2 (4DGP[1]) and from the crystal structure of open SHP2$^{E76K}$ (6CRF[3]). The distributions of the RMSD from 4DGP and from 6CRF are reported as marginal plots.

**Movie S1.** Typical trajectory of the roto-translation of the C-SH2 domain relative to PTP in the truncated SHP2 ΔN-SH2, as described by principal components PC1 and PC2. Principal components were calculated from the rigid core of the C-SH2 domain after least-square fitting over the core of the PTP domain. The C-SH2 domain is colored in orange, the PTP domain in pink.

**Movie S2.** Exemplary structures derived from the solution ensemble of the constitutively active $SHP2^{E76K}$ mutant. The solution ensemble of $SHP2^{E76K}$ comprises a set of conformational arrangements where the N-SH2 domain is either close to the surface of the PTP domain or completely detached from PTP.

**Movie S3.** Small-angle X-ray scattering (SAXS) curve calculated from the MD ensembles of the first 500 most populated clusters of $SHP2^{E76K}$ (red line). For comparison, the experimental curve (black dots), reported as raw data, and the single-structure model by Pádua *et al.* (green line)[2] are shown. Radii of gyration ($R_g$) from the MD ensembles and from experiment are reported with black and red font, respectively. The $R_g$ values from MD ensembles were obtained by Guinier fits to calculated SAXS curves, thus the values include contributions from the hydration layer.
